# A novel nabelschnur protein regulates segregation of the kinetoplast DNA in *Trypanosoma brucei*

**DOI:** 10.1101/2024.03.18.585547

**Authors:** Lawrence Rudy Cadena, Michael Hammond, Martina Tesařová, Ľubomíra Chmelová, Michaela Svobodová, Ignacio M. Durante, Vyacheslav Yurchenko, Julius Lukeš

**Affiliations:** Institute of Parasitology, Biology Centre, Czech Academy of Sciences, České Budějovice (Budweis), Czech Republic; Faculty of Sciences, University of South Bohemia, České Budějovice (Budweis), Czech Republic; Life Science Research Centre, Faculty of Science, University of Ostrava, Ostrava, Czech Republic

## Abstract

The kinetoplast DNA (kDNA), a distinctive arrangement of mitochondrial DNA found in trypanosomatid protists, comprises a concatenated network of minicircles and maxicircles that undergo division and segregation once during each cell cycle. Despite the identification and characterization of numerous proteins involved in kDNA maintenance and replication, its segregation and the formation of the nabelschnur remain poorly understood on a molecular level. This enigmatic filamentous structure, transiently appearing in *Trypanosoma brucei,* connects the daughter kDNA networks prior to their complete segregation. Here, we characterize TbNAB70, a high mobility group box-like protein localized exclusively to the nabelschnur and the kDNA disc. Our findings demonstrate that TbNAB70 is critical for the segregation, but not replication, of kDNA, a so far unprecedented phenotype. Furthermore, structural predictions suggest that this protein holds the capacity to bind to kDNA illuminating the exact molecular mechanisms of segregation involved. Thus, we propose that TbNAB70 plays a pivotal role in the faithful and efficient segregation of the daughter kDNA networks.

## INTRODUCTION

The acquisition of mitochondria was imperative for initiating eukaryogenesis and thus is a characteristic feature of eukaryotic cells (1–4). This organelle performs a multitude of different functions ranging from catabolic reactions, such as oxidative phosphorylation and fatty acid oxidation (5), to anabolic processes, namely iron-sulfur cluster assembly (6) and Ca^2+^ homeostasis (7). While the vast majority of mitochondrial proteins are encoded by the nuclear genome, a small set of proteins, mostly those of respiratory chain complexes, remain encoded by the organellar genome (8,9).

The parasitic protist *Trypanosoma brucei* contains a singular mitochondrion with a unique mitochondrial genome, termed the kinetoplast DNA (kDNA) (10). This complex structure is one of the hallmarks of the family Trypanosomatidae, a species-rich group of parasitic flagellates encompassing various causative agents of serious human diseases, such as *Trypanosoma cruzi* and *Leishmania* spp. that along with mostly free-living Bodonidae forms the order Kinetoplastea (11). In all trypanosomatids, kDNA consists of two categories of non-supercoiled open circular DNA molecules known as maxicircles and minicircles (12). In *T. brucei*, the 23 kb-long maxicircle is present in ~25 copies and codes for 18 proteins and 2 mitochondrial ribosomal RNAs (13,14), 12 of which are encoded by cryptogenes, meaning that their primary transcripts must be converted into translatable mRNA by RNA editing, a process consisting of multiple post-transcriptional uridine insertions and deletions (15,16). The ~7,500 copies constituting approximately 5,000 unique ~1.0 kb-long minicircles encode hundreds of small guide RNAs that mediate RNA editing of most mitochondrial transcripts (17–19). All minicircles are catenated into a single network forming an ultrastructurally prominent disc-shaped structure that constitutes > 90% of the kDNA mass (20,21). The maxicircles are interwoven into the minicircle network and additionally are interlocked with each other (10).

Replication of the kDNA occurs during the G_1_ phase of the cell cycle, prior to the start of nuclear DNA replication (22). A current model for kDNA replication assumes that at its initiation, minicircles are released into the kinetoflagellar zone (KFZ), where they undergo replication (23,24). The replication products are then separated and transported by a yet-to-be-described mechanism to the opposing ends of the kDNA disc, known as the antipodal sites (APS), where they are processed further and eventually re-attached to the network (23,25,26). Once the minicircles have been replicated and reattached, the remaining nicks and gaps are repaired, allowing the replication machinery to distinguish between the already replicated (i.e. nicked, gapped) and still non-replicated (i.e. covalently closed) minicircles, ensuring that each minicircle is replicated only once per generation (27,28). Similarly to minicircles, maxicircles replicate uni-directionally *via* the θ intermediates, although they remain attached to the network throughout this process (29). The segregation of the daughter kDNA networks is mediated by separation of the basal bodies of the flagellum (30). The unique physical connection between the kDNA and the basal bodies affecting this segregation is a filamentous structure transferring both mitochondrial membranes and is termed the tripartite attachment complex (TAC) (30). This structure is subdivided into three regions: (1) the exclusion zone filaments, located between the basal bodies and the outer mitochondrial membrane that is devoid of ribosomes, (2) the differentiated mitochondrial membranes, and (3) the unilateral filaments that connect the inner mitochondrial membrane to the kDNA (20,31).

While numerous proteins found within the kDNA, TAC, KFZ, and APS have been functionally characterized and identified as vital components of kDNA replication and division, the molecular mechanisms governing this highly precise process continues to be largely unknown (20,26). One division-related and morphologically characteristic structure that remains most enigmatic is the “nabelschnur”, an undefined, non-nucleic acid, filament-resembling structure observed by electron microscopy between segregating daughter kDNA networks (32,33). This structure, termed ‘nabelschnur’ (umbilicullus in German) has thus far been observed exclusively in *T. brucei*, most likely due to it being the model trypanosomatid, but is plausible to be found in other trypanosomatids as well. It has been proposed to constitute the final physical connection between the newly replicated kDNA networks (10). To date, only one protein, TbLAP1 (Tb927.8.3060), a M17 family leucyl aminopeptidase metalloprotease is known to localize to the nabelschnur, as well as to the APS (34). Knockdown of TbLAP1 leads to growth arrest prior to cytokinesis, while its overexpression results in a loss of kDNA, suggesting a role in the kDNA segregation. The TbLAP1 knockdown phenotype is ultimately not lethal for cells, and kDNA is observed successfully segregating, albeit at a slower rate, suggesting the involvement of at least one other unknown component regulating this process.

While screening proteins from the *T. brucei* MitoTag project (35), we identified a previously uncharacterized protein with an mNeonGreen-signal appearing to localize to the kDNA as well as forming a point of connection between certain dividing kDNA. In this work, we demonstrate that this kDNA-associated protein, named TbNAB70, indeed localizes to the nabelschnur and plays an essential role in the segregation of newly replicated kDNAs and subsequent cytokinesis in *T. brucei*.

## RESULTS

### TbNAB70 localizes to the kDNA and nabelschnur, exhibiting cell cycle specific expression

TbNAB70 (Tb927.11.7580) is a basic (pI = 9.18), 70 kDa hypothetical conserved protein with a predicted mitochondrial targeting sequence at its N-terminus. When screening for proteins exhibiting a distinct kinetoplast signal in frame of the MitoTag project (35), TbNAB70 had been identified as an uninvestigated protein that warranted further scrutiny.

To localize TbNAB70 and verify whether it represented a genuine component of the kDNA, we endogenously tagged its C-terminus with a V5_×3_ epitope in SmOx procyclic form *T. brucei* (36,37). Colocalization analysis with DAPI confirmed that TbNAB70 localizes to the kDNA (Fig. 1A; Fig. S1). Its positioning during early phases of the cell cycle varied from the kDNA in the G_1_ phase to a dumbbell-like structure co-localizing with the kDNA disc mimicking this elongated structure in early 1Kdiv1N cells (Fig. 1A). Shortly afterwards, the discs commence segregation into the daughter kDNA networks positioned perpendicular to one another.

**Fig. 1.**
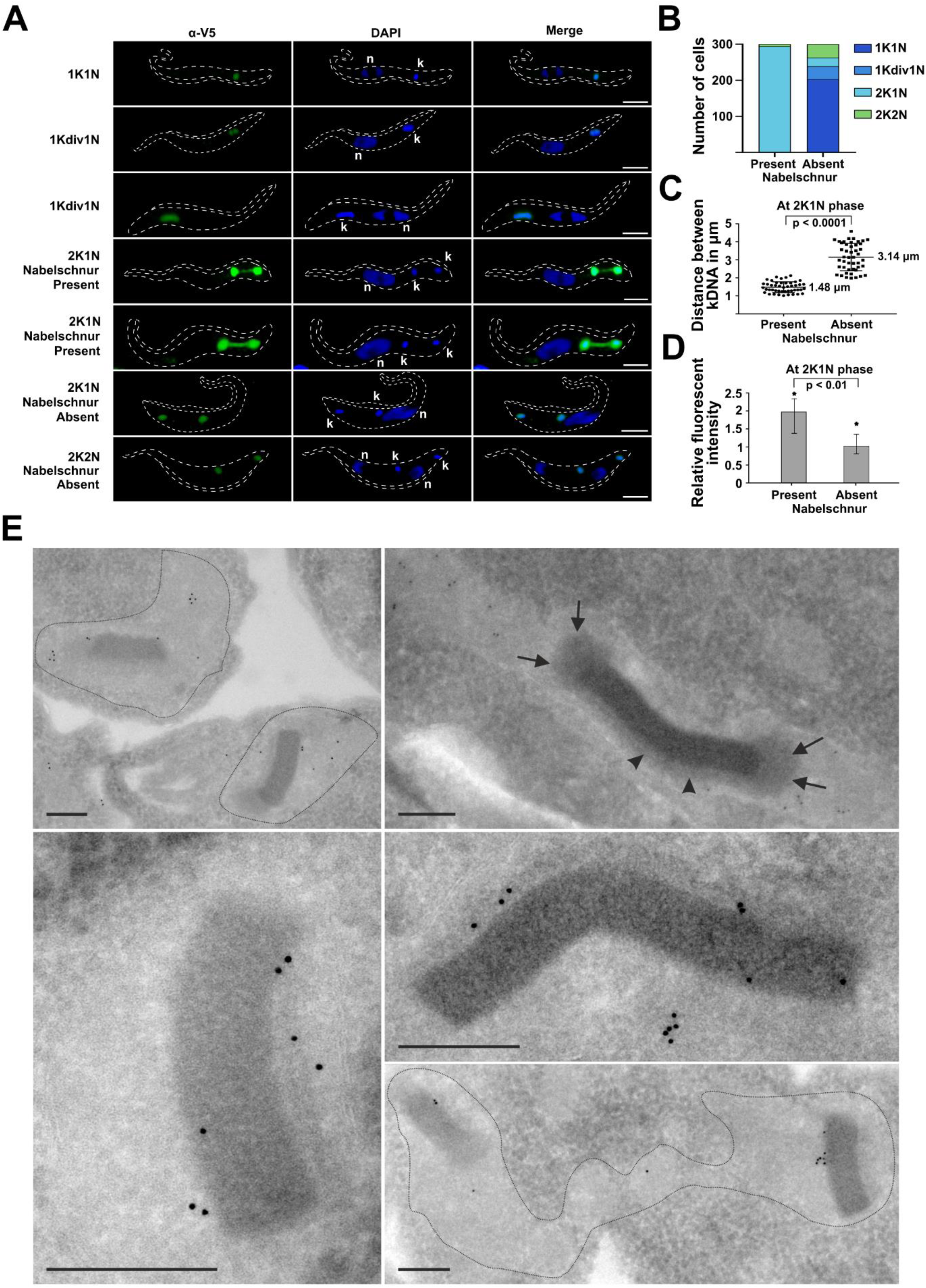
Localization of TbNAB70 in procyclic *T. brucei*. (**A**) Representative immunofluorescence microscopy images of TbNAB70_V5_ procyclic cells during different stages of the cell cycle (1K1N, 1Kdiv1N, 2K1N, 2K2N). TbNAB70_V5_ (green) was detected by means of α-V5 antibody. The kDNA (k) and the nucleus (n) were stained with DAPI (blue). Scale bars: 2 μm. (**B**) Graph depicting presence and absence of TbNAB70_V5_-visualized nabelschnur during specific stages of cell cycle. 100 cells were counted at random with three biological replicates (n=3), resulting in a total of 300 cells represented on the graph. (**C**) Quantitative analysis of distance between two daughter kDNA networks during 2K1N phase measuring length when the nabelschnur disappears. (**D**) Relative fluorescent intensity of TbNAB70_V5_ signal associated to the kDNA during 2K1N phase while the nabelschnur is present or absent. Forty cells with nabelschnur and 40 cells without nabelschnur were measured with the average intensity of green fluorescence seen in the nabelschnur-lacking 2K1N cells set to 1 as a value. Asterisks show maximum value. (**E**) Transmission electron microscopy of cryosections of TbNAB70_V5_ cells immunolabelled with Protein A 10 nm. Arrows and arrowheads point towards the antipodal sites and the kinetoflagellar zone of the kDNA, respectively. Scale bar: 250 nm.

At this stage, the characteristic nabelschnur between the newly replicated kDNA networks becomes apparent. As the daughter kDNA discs progressively separate (2K1N stage, note the apparent lack of the DAPI signal in the space between them), the nabelschnur remains continually present (Fig. 1A-B). This structure begins to disappear and disintegrate only at distances greater than approximately 2 μm between the daughter kDNA networks during the 2K1N stage, when the TbNAB70 signal remains confined to two regions overlaying each newly synthesized kDNA disc only (Fig. 1C). Noteworthy, the fluorescent intensity of the tagged TbNAB70_V5_ increases when the nabelschnur is present and diminishes once the structure disappears at the 2K1N phase (Fig. 1D) strongly suggesting that the level and distribution of TbNAB70 are strictly stage-specific. In order to explicitly verify the localization of TbNAB70_V5_, we employed immunodecoration of cryosectioned cells processed for transmission electron microscopy using gold-labeled α-V5 antibody. We documented numerous gold particles on the periphery of the electron-dense kDNA disc in addition to areas where the presumed nabelschnur is localized (Fig. 1E). Therefore, we conclude that TbNAB70 dually localizes to the nabelschnur and the kDNA periphery, and its expression is cell cycle-specific.

### Depletion of TbNAB70 by RNAi demonstrates an essential function in kDNA segregation

To study the function of TbNAB70, we depleted the corresponding mRNA by RNA interference using a tetracycline-inducible RNAi vector (38) in the TbNAB70_V5_ SmOx procyclic trypanosomes, as described above. The RNAi knockdown was efficient, as judged by probing for the TbNAB70_V5_ protein in western blots that showed its complete disappearance by 48 hours post-RNAi induction (Fig. 2A). The cells displayed a severe growth defect within 24 hours post-RNAi induction, indicating that this protein is critical for cell viability (Fig. 2B). Intrigued by the severity of the growth defect, we performed Mitotracker Red staining of cells 48 hours post-RNAi induction and witnessed neither gross morphological defects of the mitochondrion, nor the loss of its membrane potential (Fig. 2C). In order to investigate whether cytokinesis was affected, we performed DAPI counts on cells before treatment, as well as 24 and 48 hours post-RNAi induction. In the non-induced cells, we observed approximately 65% of cells exhibiting 1K1N (G_1_ phase) phenotype. That ratio decreased to 53% 24 hours post-RNAi induction and dropped further down to 40% another 24 hours later (Fig. 2D). Additionally, we recorded that ~15% of cells depicted a 1K2N phenotype at 24 hours, with an increase to ~25% at 48 hours post-RNAi induction. This 1K2N phenotype was not observed in any of the non-induced cells counted indicating that both the kDNA segregation and cell cycle were affected upon depletion of TbNAB70. Interestingly, 10% of cells showed 1Kdiv2N 24 hours post-RNAi induction, although within next 24 hours this decreased to just 5%. Moreover, the same decrease was observed for cells exhibiting 2K1N and 2K2N phenotype.

**Fig. 2.**
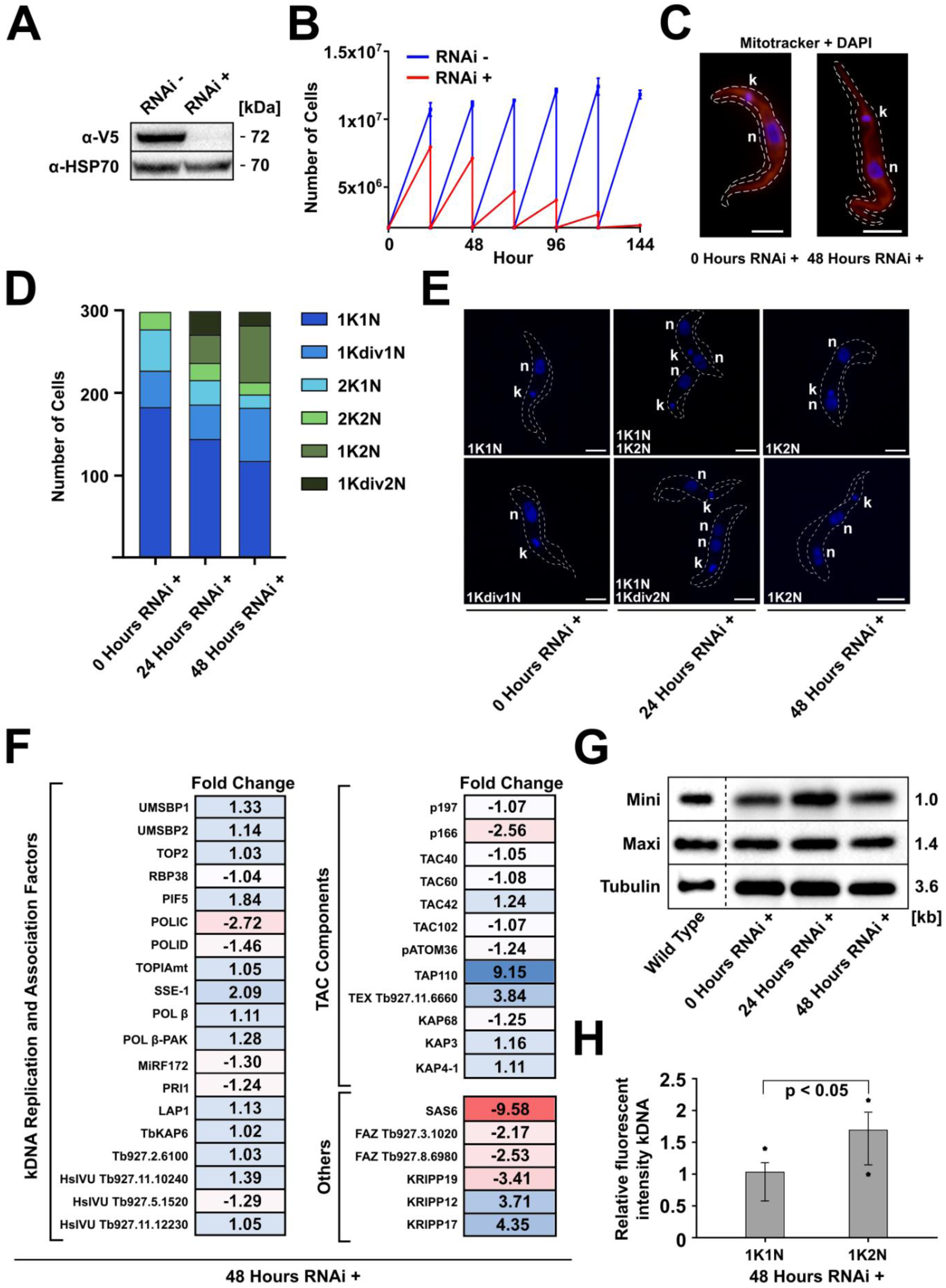
Depletion of TbNAB70 mRNA by RNAi. (**A**) Western blot of whole cell lysates showing complete depletion of TbNAB70_V5_ at 48 hours post-RNAi induction. TbNAB70_V5_ was detected by α-V5 antibody. Heat shock protein 70 (HSP70) served as a loading control. (**B**) Growth curve of non-induced and RNAi-induced TbNAB70_V5_ cells (n = 3, mean and standard deviation values are shown). (**C**) Immunofluorescence images of non-induced cells (0 hours RNAi +) and induced cells (48 hours RNAi +) stained with Mitotracker Red (red), with kDNA networks (k) and nuclei (n) as detected as in Fig. 1. Scale bar: 2 μm. (**D**) Quantification of the relative occurrence of kDNA networks and nuclei in the non-induced cells (0 hours RNAi +), and induced cells (24 hours RNAi + and 48 hours RNAi +). 100 cells were chosen at random and analyzed. Three biological replicates were performed (n = 3) resulting in a total of 300 cells depicted on the graph. Mean and standard deviation values are shown. (**E**) Representative immunofluorescence images of non-induced cells (0 hours RNAi +), and induced cells (24 hours RNAi + and 48 hours RNAi +) depict kDNA networks (k) and nuclei (n) as quantified in (D). (**F**) Heatmap showing increase or decrease fold change in selected proteins encompassing kDNA replication and associated factors, TAC components, and other proteins of interest in cells 48 hours post-induction. For the full list of proteins, see Suppl. Table 1. (**G**) Southern blot analysis of wild type, non-induced cells (0 hours RNAi +), and induced cells (24 hours RNAi + and 48 hours RNAi +). Numbers indicate size of DNA fragments in kb. (**H**) Relative fluorescent intensity of DAPI stained kDNA in 48-hour RNAi-induced cells. kDNA intensity was measured in twenty cells depicting the phenotype 1K1N and compared kDNA intensity of twenty cells depicting the phenotype 1K2N. The average intensity of kDNA in the 1K1N cells was normalized to 1 as a set value. Asterisks show maximum and minimum values. Mean and standard deviation data are shown.

To test whether TbNAB70 depletion had an effect on the components of TAC or other kinetoplast-associated proteins, we performed quantitative proteomics comparing the non-induced cells and those 48 hours post-RNAi induction in triplicates (Fig. 2F). We observed no significant changes in protein levels of the cohort of kDNA replication and associated factors, save for a −2.72 fold decrease in levels of POLIC, a protein suggested to play a role in maxicircle replication that dually localizes to the KFZ and APS (39,40). It is worth noting that three components of the TAC also displayed significant changes in their expression levels, most prominently TAP110 (Tb927.11.7590) and Tb927.11.6660, which were both upregulated 9.15 and 3.84 times, respectively. Such an increase of these two proteins (in comparison to other TAC components) is of particular interest, as it has been reported that overexpression of TAP110 led to a strong increase of Tb927.11.6660, indicating these two proteins may form a yet undescribed interaction with each other. Furthermore, TAP110, known to play a supporting role in the separation of replicated kDNA networks (41), is present directly upstream of TbNAB70 within the genome of *T. brucei*. Simultaneously, there was a decrease in the level of p166, another characterized TAC component of the mitochondrional inner membrane (42). We chose to not restrict ourselves in quantifying proteins known to associate to the kDNA or TAC; noting a strong decrease (−9.58-fold reduction) of the basal body protein SAS6 and, to a lesser degree, components of the flagellar attachment zone (FAZ). Additionally, we witnessed significantly reduced or upregulated levels of numerous large and small subunits of the mitochondrial ribosome and KRIPP proteins (Fig. 2F, Table S1).

Next, we decided to verify if TbNAB70 depletion leads to changes in the abundance of minicircles and/or maxicircles potentially indicating whether there is an effect on the kDNA replication. We employed Southern blot analysis of total DNA isolated from trypanosomes 0, 24 and 48 hours post-RNAi induction, in addition to wild type trypanosomes, using specific probes against minicircles and maxicircles, with tubulin serving as a loading control. As shown in Fig. 2G, no changes in the levels of both monitored kDNA components occurred following the TbNAB70 depletion. This lack of change in gross abundance suggests that kDNA replication itself is not affected when normalized to total DNA amount in a pool of cells during different cell cycle stages. However, this method cannot distinguish whether kDNA replication has taken place in cells where one kDNA is morphologically present alongside two nuclei (1K2N phenotype). To verify if the kDNA has indeed replicated, yet not separated in cells during this stage, we quantified the DAPI intensity of the kDNA in individual cells showcasing the 1K1N phenotype and compared it to that found within the 1K2N cells. We measured a near 2-fold increase in the kDNA intensity in the 1K2N cells 48 hours post-RNAi induction (Fig. 2H), indicating that the kDNA content has nearly doubled yet molecules had not segregated.

### TbNAB70 depletion ultimately leads to inhibition of flagellogenesis-related processes

Proteomics data of TbNAB70 RNAi cells suggested prominent depletion of the FAZ and basal body components (Fig. 2F). Accordingly, we sought to visualize observable changes in these regions using antibodies against FAZ1 (Tb927.4.3740) and Y1/2, the latter being a marker for the mature basal body of *T. brucei*. Our observations of 1K2N cells appearing 48 hours post-RNAi induction show the absence of a second mature basal body suggesting that cells are unable to generate this structure following the NAB70 ablation (Fig. 3). Furthermore, such 1K2N cells display a single FAZ signal, indicative of an inability to synthesize a second FAZ. Some of these cells additionally display an extremely reduced singular FAZ signal with a detached flagellum accompanied by aberrant cell morphology (Fig. 3).

**Fig. 3.**
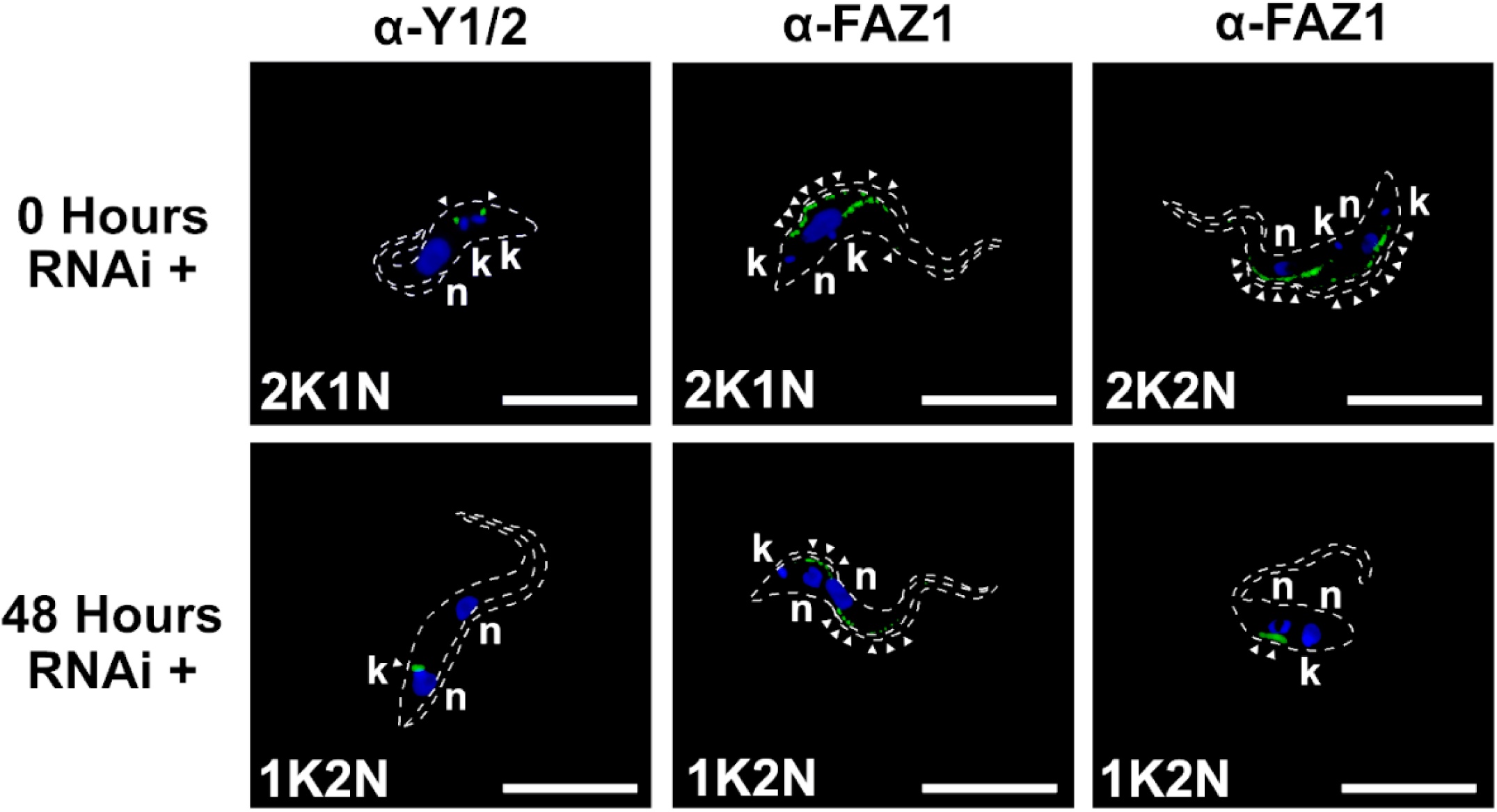
Loss of TbNAB70 leads to flagellogenesis inability. Immunofluorescence images of non-induced cells (0 hours RNAi +) and induced cells (48 hours RNAi +). Mature basal body stained with α-Y1/2 (green – first panel) or flagellum stained with α-FAZ1 (green – second and third panel), with kDNA networks (k) and nuclei (n) detected as in Fig. 1. Arrowheads indicate α-Y1/2 or α-FAZ1. Scale bar: 10 μm. Note the detachment of the flagellum on the bottom right image.

### *In silico* structural analysis of TbNAB70 indicates a capacity for physical kDNA binding

Observing the localization of TbNAB70 and effects of its RNAi knockdown on post-replicated kDNA segregation, we ventured further in elucidating the functionality on a molecular level. *In silico* analysis using SignalP revealed that TbNAB70 contains a 23 amino acid-long mitochondrial targeting sequence at the N-terminus. Additionally, hits for a coiled coil domain between amino acid positions 278-306 and an intrinsically disordered domain at the C-terminus were revealed (Fig. 4A). By performing a HHpred search, we further analyzed whether TbNAB70 has any homologous hits or regions to other previously characterized proteins. While no homologous proteins encompassing the entirety of TbNAB70 were detected, one particular region (amino acids 382-484) showed a strong homology to various mitochondrial DNA-binding proteins (Table S2), such as Gcf1p of *Candida albicans* containing High Mobility Groups (HMG) box domains (43).

**Fig. 4.**
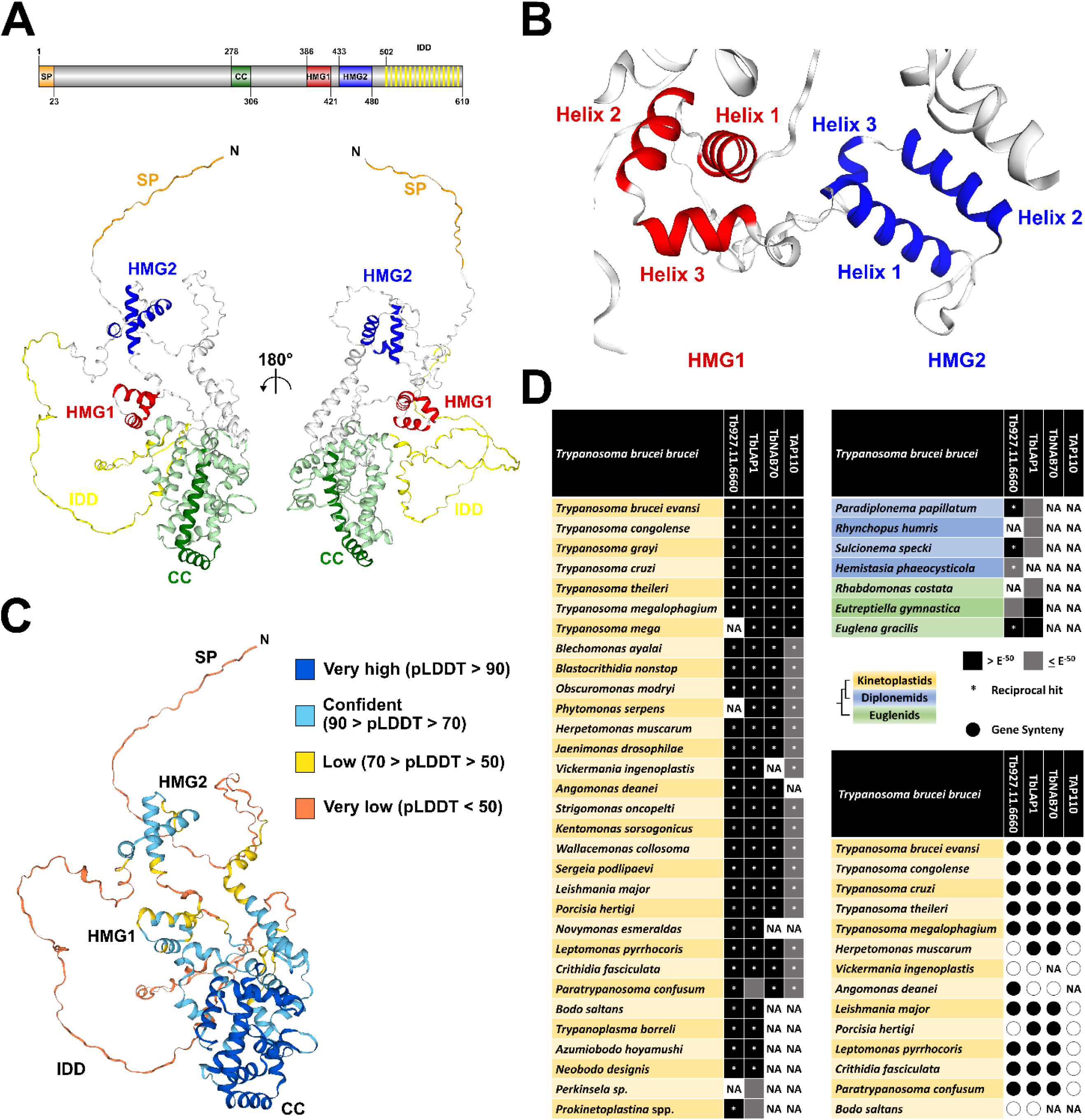
*In silico* analysis of TbNAB70 and phylogeny of associated proteins. (**A**) Alphafold2 structure of TbNAB70. The predicted signal peptide (SP) is in orange, the coiled coil domain (CC) in dark green, regions of highest confidence in light green, High Mobility Group box 1 (HMG1) in red, High Mobility Group box 2 (HMG2) in blue, and predicted intrinsic disordered domain (IDD) in yellow. The linear map of the domains is shown above. (**B**) Close up view of tertiary structure of HMG box 1 and HMG box 2. (**C**) Gross confidence scores from Alphafold2. (**D**) Prescence and conservation of TbNAB70 and relevant genes (TbLAP1, TAP110 and TEX like-Tb927.11.6660) across the Euglenozoan phylum as well as an assessment of synteny across high quality genomes of Trypanosomatidae, sequence information available in Table S3.

Next, we predicted the tertiary structure of TbNAB70 using AlphaFold2 (Fig. 4A-C), which revealed the presence of three small α helices confined in a triangular space between amino acid positions 386-421 and another three small α helices in a similar triangular space between positions 433-480 (Fig. 4A-B). Since all these motifs are hallmarks of the HMG box domains (44,45), we term them HMG boxes 1 and 2. The AlphaFold2 results show a confident prediction of these two HMG domain regions (90 > pLDDT > 70) (Fig. 4C). Additionally, the coiled coil domain showed the highest confidence (pLDDT 90 >) while the predicted intrinsically disordered region at the C-terminus showed a very low confidence score (pLDDT < 50), a typical score for predicted disordered regions (46).

### *TbNAB70* is restricted to parasitic kinetoplastids and likely developed in tandem with *TAP110*

We decided to investigate the phylogenetic distribution of *TbNAB70* within a spread of available genomes and transcriptomes for representatives of the phylum Euglenozoa in comparison to the only other characterized nabelschnur protein, TbLAP1 (Fig. 4D) (34). Similarly, we opted to assess the TAC components TAP110 and TEX-like protein encoded by *Tb927.11.6660*; which displayed notable enrichment following TbNAB70 RNAi induction.

Of the two identified nabelschnur components, *TbLAP1* shows conserved distribution in all surveyed kinetoplastid flagellates, with homologues additionally present in other Euglenozoa and likely beyond, suggesting an emergence that preceded the development of the kinetoplast structure itself (Fig. 4D). Additionally, several M17 metalloproteases are present within the genome of *T. brucei*, and accordingly, reciprocal hits to *TbLAP1* were not produced in euglenids, diplonemids and certain kinetoplastid members more distantly related to *T. brucei* (Fig. 4D). Conversely, *TbNAB70* is confined to Trypanosomatidae, a clade of exclusively parasitic kinetoplastids. Co-enrichment of the TAC proteins Tb927.11.6660 and TAP110 suggests putative interaction or supporting roles for these two proteins. Despite this, these genes show contrasting levels of distribution, with TEX-like *Tb927.11.6660* seen in the majority of Euglenozoa, while *TAP110* shows a highly complementary distribution to that of *TbNAB70*, suggesting a similar time of emergence for these two proteins. Shared synteny is seen in all four proteins throughout the genus *Trypanosoma* (47), while additionally being broadly present in the clade of Leishmaniinae as well as in the early-branching *Paratrypanosoma confusum,* except for *TAP110* (Fig. 4D). Such a discrepancy is notable considering that *TAP110* and *TbNAB70* are neighboring genes, suggesting a break in this syntenic region occurring immediately upstream to *TAP110* in the common ancestor of this clade. *TbNAB70* is absent from *Vickermania ingenoplastis* and shows a lack of synteny for all other surveyed genes in this study, indicating a high level of gene movement and disruption in this organism. *TAP110* has additionally been lost in *Angomonas deanei*, while both genes are absent from *Novymonas esmeraldas*, which prompts speculation on these organisms’ execution of kDNA division.

## DISCUSSION

Until now, TbLAP1 represented the only identified protein that localized to the nabelschnur, a filamentous structure that forms the physical connection between two dividing daughter kDNA networks during cytokinesis of *T. brucei* (34). Here, we report the identification of TbNAB70, a second protein localizing to the nabelschnur, as well as the kDNA disc periphery. Despite being a basic protein, TbNAB70 interacts not only with the kDNA, but also with the non-nucleic acid filaments that constitute the nabelschnur. Moreover, we show that the expression of TbNAB70 is cell cycle-specific, demonstrating greater abundance in cells during kDNA division (2K1N) compared to those in G_1_.

Knockdown of TbLAP1 has been shown to result in an accumulation of 2K2N cells, delaying the completion of cytokinesis and with a reduced growth rate, but proving nonlethal (34). By contrast, TbNAB70 displayed a drastic growth defect accompanied by a significant increase of 1K2N cells with replicated kDNA, that is ultimately unable to segregate. Furthermore, the cells fail to build a mature second basal body, and, in some cases, display a complete disattachment of the mother flagellum. The nabelschnur represents the linker between two replicated kDNAs, with its process of dissolution or breakdown yet undetermined. Although TbLAP1 contains a metalloprotease domain, its ablation suggests that this protein does not participate in cleaving of the nabelschnur, evidenced by successfully divided kDNA discs. By contrast, TbNAB70 appears to be functionally active immediately prior to kDNA segregation and thus represents a singular candidate to be regulating this dissolution step.

Depletion of TbNAB70 does not lead to dramatic changes in abundance of known components of the kDNA replication machinery as well as most TAC complex, although two of these proteins, TAP110 and Tb927.11.6660 (a TEX-like protein), showed significant upregulation. It was previously demonstrated that an overexpression of TAP110 leads to an upregulation of Tb927.11.6660 (41) indicating that these two proteins may interact in a yet uncharacterized manner. Although it has been proposed that TAP110 is a kDNA segregation factor, with Tb927.11.6660 potentially providing a link between mitochondrial-nuclear communication (41), neither localize to the nabelschnur, indicating that TbNAB70 plays a greater role in kDNA segregation than these other kDNA-associated proteins. We suggest that increases in TAP110 levels, following TbNAB70 depletion, represents a compensatory mechanism responding to segregation difficulties encountered post-replication of kDNA, which is supported by the neighboring positions of these genes, combined with their putative annotation. Unfortunately, in the aforementioned study of TAP110 overexpression analysis we were unable to identify whether TbNAB70 levels were affected, as the mass spectrometry dataset is not publicly available (41).

Leucine amino peptidases are observed in a wide variety of organisms and they perform a diverse range of functions (34,48). Based on their phylogenetic distribution, *TbLAP1* and *Tb927.11.6660* preceded the development of the kDNA, and likely represent proteins repurposed or adapted to mediate processes associated with this unique structure. By contrast, *TbNAB70* and *TAP110* superseded the emergence of the kDNA network, itself a complex and evolutionary improbably structure (49). These two proteins seem to have appeared in tandem, as suggested by their neighboring genomic position as well as their distribution in kinetoplastids. We speculate that these proteins may have fulfilled the functional requirements of an increasingly complex organelle substructure, that already is known to mandate dozens of proteins for its maintenance (10,17). Moreover, our knockdown analysis indicates that in terms of successful kDNA segregation and cell cycle completion, TbNAB70 may have rendered TbLAP1 partially redundant. Based on our survey of gene conservation, *Novymonas esmeraldas* represents the ideal organism to investigate this hypothesis, having lost both ‘recent’ additions of *TbNAB70* and *TAP110*, combined with its recently established tractability to investigate specific genes of interest (50,51). Coincidently, we also note that two of three surveyed organisms which have specifically lost *TbNAB70* and/or *TAP110*, also bear bacterial endosymbionts, endosymbiotic acquisitions representing a comparatively rare occurrence among trypanosomatids (52). Whether a causative relationship exists between these two events remains an exciting possibility to be explored.

Structural predictions of TbNAB70 identified a coiled coil domain and an intrinsic disordered domain spanning the C-terminus. Although this protein is confined to the obligatory parasitic kinetoplastids, using Hidden Markov Model analysis, we have found hits to various mitochondrial DNA binding proteins containing high mobility group boxes, domains known to interact with high affinity to non-B type DNA, such as kinked or unwounded DNA during the replication process (44), with the top hit being the HMG box protein Gcf1p with an experimentally solved structure (43,53). The predicted tertiary structure of TbNAB70 revealed two regions containing 3 small α-helices confined in a triangular space, hallmarks of high mobility group box domains (44,45), allowing us to conclude that TbNAB70 belongs to this protein category. It should be noted, however, that TbNAB70 is not the only HMG-box-containing protein localizing to the kDNA, as the previously described TbKAP6 has been shown to contain a single HMG-box domain. Functional studies using RNAi targeting TbKAP6 resulted in a loss of minicircles and maxicircles, while its overexpression led to an increase of decatenated minicircles (54). This supports the idea that TbKAP6 is involved in the minicircle release process prior to replication, but not kDNA segregation, as is the case with TbNAB70. Although both proteins contain HMG-box domains (44), they do not appear to directly interact with each other based on our depletome analysis and TbNAB70’s unique localization to the nabelschnur.

In summary, we assume that during and/or after kDNA replication, TbNAB70 interacts with the kDNA mediating the separation of the daughter networks, possibly through movements of the intrinsic disordered domains, forming the nabelschnur in the process. Following the depletion of TbNAB70, the basal bodies are unable to segregate in the absence of divided kDNA and, as a result, the levels of FAZ components and SAS6, a protein associated with the biogenesis of probasal bodies (55), are decreased and cytokinesis is ultimately arrested (Fig. 5). Although our findings strongly support the view of a nabelschnur-mediated kDNA segregation mechanism, the exact composition of this enigmatic structure is still unknown. Moreover, whether TbNAB70 indeed physically interacts with the kDNA, and whether this interaction is specific to the maxicircles or minicircles, remains to be investigated.

**Fig. 5.**
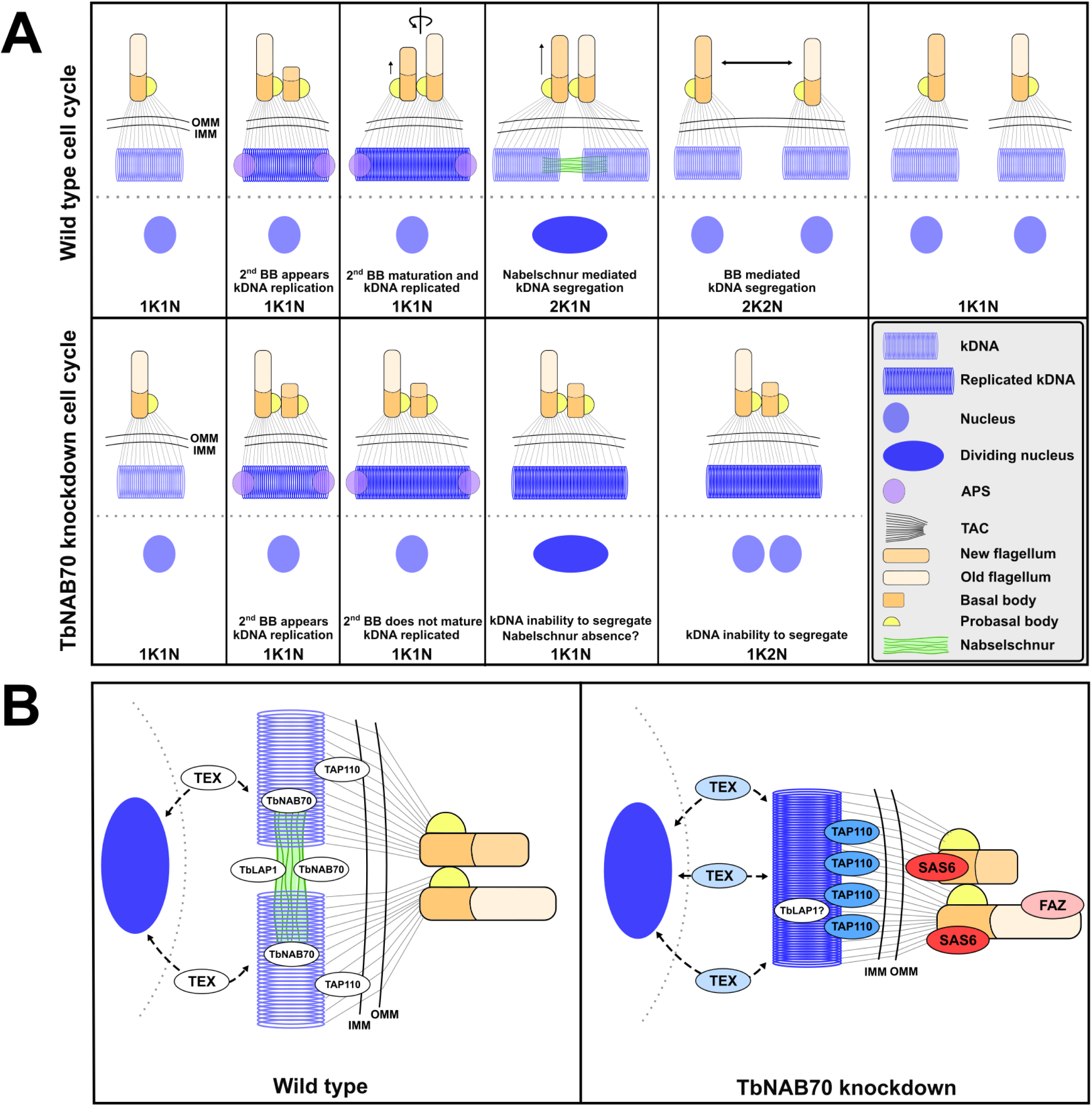
Mechanism of action model of TbNAB70 dependent kDNA segregation. (**A**) Model of the proposed mechanism of replicated daughter kDNA segregation throughout the *T. brucei* cell cycle. OMM marks the outer mitochondrial membrane and IMM is the inner mitochondrial membrane. **(B)** Model of interactions of selected kinetoplast and flagellum-associated proteins.

In conclusion, the identification of TbNAB70 sheds light on the mechanisms of kDNA segregation, and thus fulfils the criteria for a long sought-after regulatory component of the nabelschnur, critical for the successful partitioning of kDNA.

## MATERIALS AND METHODS

### Generation of *T. brucei* transgenic cell lines

*Trypanosoma brucei* SmOxP927 procyclic cell line (36,37) was transfected by electroporation using the Amaxa apparatus (Lonza) set with the program X-14. After transfection, cells were seeded in 24-well plates in serial dilutions and selected by the appropriate resistance marker. For endogenous V5_×3_ epitope, a previously described strategy was employed (37). Briefly, C-terminus endogenous tagging amplicons were obtained using long primers targeting a modified pPOTv4 vector containing a V5_×3_ epitope. Cells were selected with Hygromycin (Thermo Fisher Scientific) at 50 μg/ml and screened for V5 expression by western blotting. For generation of the RNAi cell line, PCR products targeting a specific CDC region as determined by RNAit (56) were cloned into the p2T7-177 vector (57). *Not*I-linearized construct was electroporated using the same conditions as above into the V5_×3_ tagged cell line. Cells were selected with Phleomycin (Sigma-Aldrich / Merck) at 2.5 μg/ml. Knockdown of the protein was monitored by western blotting against the V5_×3_ epitope using α-V5 mouse primary antibody (Invitrogen / Thermo Fisher Scientific – 1:1000). Oligonucleotides used for PCR amplification are given in Table S4.

### Cell culturing

Cells were grown at 27°C in SDM-79 medium supplemented with 10 % fetal bovine serum and 7.5 mg/l hemin. Cells were grown in the presence and absence of 1 μg/ml tetracycline (Sigma-Aldrich / Merck) for RNAi induction. Cell density was measured using the Beckman Coulter Z2 cell and particle counter (Beckman Coulter) and maintained at the exponential mid-log growth phase throughout the analysis.

### Immunofluorescence microscopy

A total of 10^6^ *T. brucei* cells were fixed with 4 % paraformaldehyde for 30 min at room temperature (RT). The cells were subsequently centrifuged at 1,300 × g for 5 min, resuspended in 200 μl 1 × PBS, and placed on a glass slide for 30 min. The slides were washed with PBS and allowed to permeabilize in methanol for 30 min at −20°C. The slides were washed thrice with PBS and blocked with 5 % milk inside a humid chamber for 1 h before being incubated with α-V5 mouse primary antibody (Thermo Fisher Scientific – 1:400) overnight at 4°C. The slides were washed once again with PBS and incubated with secondary Alexa Fluor 488 goat α-mouse IgG antibody (Invitrogen / Thermo Fisher Scientific – 1:1,000) for 1 h at RT. The slides were washed with PBS before a final wash with water, air-dried, and mounted in ProLong Gold antifade reagent DAPI (Invitrogen / Thermo Fisher Scientific). Images were captured with the BX51 Fluorescence Microscope (Olympus). For Mitotracker Red staining (Invitrogen / Thermo Fisher Scientific), the same procedure was performed with the exception of incubating cells in culture with the staining 30 min prior to fixation. Fluorescent intensity analysis was performed using ImageJ (58).

Mature basal bodies were illuminated as above with monoclonal rat antibody Y1/2, specific to tyrosinated tubulin (1:500) (59) followed by secondary Alexa Fluor 488 goat α-mouse IgG antibody (Invitrogen / Thermo Fisher Scientific – 1:100). FAZ was visualized via anti-FAZ1 antibody L3B2 (antigenic to Tb927.4.3740, 1:3) (60) followed by secondary Alexa Fluor 488 goat α-mouse IgG antibody (Invitrogen / Thermo Fisher Scientific – 1:1,000).

### Transmission electron microscopy and immunogold labeling

The cell pellet was fixed in a mixture of 4% formaldehyde and 0.1% glutaraldehyde in 0.1M HEPES (Sigma-Aldrich / Merck) for 1 h at RT, resuspended and washed with 0.1M HEPES, embedded into 10% gelatin (Serva Electrophoresis) and subsequently transferred into 2.3M sucrose solution for 72 h at 4 °C. Next, the pellet was cut in small gelatin blocks, which were mounted onto flat copper stubs and then plunge frozen in liquid nitrogen. Ultrathin cryosections (100 nm) were generated using a cryo-ultramicrotome (Leica) at −100 °C. The sections were picked up using a drop of 1% methyl cellulose/1.05M sucrose mixture and transferred onto a formvar-carbon coated grid. The sections were treated with blocking buffer (TEM-BB) composed of 1% fish skin gelatin (Sigma-Aldrich / Merck) dissolved in 0.1M HEPES for 1 hour, and then incubated with the primary α-V5 monoclonal antibody (Thermo Fisher – 1:30 dilution) for 30 min at RT. After washing in TEM-BB, the sections were incubated with the secondary α-protein A monoclonal antibody conjugated with gold nanoparticles (UMC Utrecht – 1:50 dilution) in TEM-BB for 30 min at RT. A control for the non-specific binding of the secondary antibody was performed by omitting the primary antibody. Finally, the cryosections were washed with HEPES, water, contrasted using a mixture of 2% methylcellulose and 3% aqueous uranyl acetate solution (9:1), and observed using a JEM 1400-Flash transmission electron microscope (JEOL) equipped with CMOS camera Xarosa (EMSIS) at accelerating voltage 120 kV.

### Western blot

Whole cell lysate from 10^6^ *T. brucei* cells were loaded onto an SDS-PAGE gel and transferred to a PVDF membrane. Membranes were probed with either anti-V5 mouse primary antibody (Invitrogen / Thermo Fisher Scientific – 1:4,000) or α-HSP70 primary antibody (see Acknowledgements – 1:10,000) and incubated overnight at 4°C. This was followed by incubation with a secondary horseradish peroxidase-conjugated α-mouse antibody (BioRad – 1:2,000) for 1 h at RT. Proteins were visualized using the Pierce ECL system (Genetica / BioRad) on a ChemiDoc instrument (BioRad).

### Southern-blot quantification of maxi- and minicircles abundance

Total DNA was isolated from mid-log phase cultures before RNAi induction and on hours 24 and 48 post RNAi induction. The cell pellets were washed in PBS and resuspended in TRI Reagent (Molecular Research Center) as per the manufacturer’s suggestions. Five μg of total DNA per lane and probe was digested with *Hind*III and *Xba*I (tubulin and maxicircles-specific probes) or *Rsa*I (minicircles-specific probes). Digested DNA was resolved on 0.75 % agarose gel and transferred to nylon membrane (Amersham). The hybridization was performed overnight using DIG Easy Hyb (Roche). Probes to detect tubulin, maxi- and minicircles were PCR amplified and labelled using PCR DIG Probe Synthesis Kit (Roche). The detection was done using DIG Luminiscent Detection Kit (Roche) following the recommendations of the manufacturer. Oligonucleotides used for PCR amplification are given in Table S4.

### Protein preparation and mass spectroscopy

Triplicate whole cell lysates homogenized by sonication in 1% SDS were processed for mass spectroscopy analysis as described previously (61). In brief, samples were resuspended in 100 mM tetraethylammonium bromide (TEAB) containing 2 % SDS. Cysteines were reduced with a final concentration of 10 mM Tris(2-carboxyethyl) phosphine hydrochloride (TCEP) and subsequently cleaved with 1 μg of trypsin overnight at 37°C. After digestion, 1 % trifluoroacetic acid (TFA) was added to wash twice, and samples were resuspended in 20 μl of TFA per 100 μg of protein. A non-reversed-phased column (Easp-Spray column, 50-cm by 75-μm inner diameter, PepMap C_18_, 2-μm particles, 100-Å pore size) was used for LC-MS analysis. Mobile phase buffer A consisted of water and 0.1 % formic acid. Mobile phase B consisted of acetonitrile and 0.1 % formic acid. Samples were loaded onto the trap column (Acclaim PepMap300 C_18_, 5 μm, 300-Å-wide pore, 300 μm by 5 mm) at a flow rate of 15 μl/min. The loading buffer consisted of water, 2 % acetonitrile, and 0.1 % TFA. Peptides were eluted using a mobile phase gradient from 2 % to 40 % over 60 min at a flow rate of 300 nl/min. The peptide cations eluted were converted to gas-phase ions via electrospray ionization and analyzed on a Thermo Orbitrap Fusion instrument (Q-OT-qIT – Thermo Fisher). Full MS spectra were acquired in the Orbitrap instrument with a mass range of 350 to 1,400 *m/z*, at a resolution of 120,000 at 200 *m/z*, and with a maximum injection time of 50 ms. Tandem MS was performed by isolation at 1.5 Th with the quadrupole, high-energy collisional dissociation (HCD) fragmentation with a normalized collision energy of 30, and rapid-scan MS analysis in the ion trap. The MS/MS ion count target was set to 10^4^, and the maximum infection time was set at 35 ms. Only those precursors with a charge state of 2 to 6 were sampled. The dynamic exclusion duration was set to 45 s with a 10-ppm tolerance around the selected precursor and its isotopes, Monoisotopic precursor selection was on with a top-speed mode of 2-s cycles.

### Analysis of mass spectrometry peptides

Label-free quantification of the data was performed using MaxQuant software (v. 1.6.2.1) (62). The false discovery rates for peptides and proteins were set to 1 % with a specified minimum peptide length of 7 amino acids. The Andromeda search engine was used for the MS/MS spectra against the *Trypanosoma brucei* database (downloaded from UniProt, December 2021, containing 8,306 entries). Enzyme specificity was set to C-terminal Arg and Lys, alongside cleavage at proline bonds, with a maximum of 2 missed cleavages. Dithiomethylation of cysteine was selected as a fixed modification, with N-terminal protein acetylation and methionine oxidation as variable modifications. The “match between runs” feature in MaxQuant was used to transfer identification to other LC-MS/MS runs based on mass and retention time with a maximum deviation of 0.7 min. Quantifications were performed using a label-free algorithm as described (62). Data analysis was performed using Perseus software (v. 1.6.1.3).

### *In silico* structural analysis

The regions homologous to different HMG box domain containing proteins was predicted using the HHpred toolkit (see Table. S2 for list of all homologous proteins with e-scores) (63). The mitochondrial targeted signal, coiled coil domain, and intrinsic disordered domain were predicted using SignalP 4.1, Coils, and MobiDB-lite, respectively (63–65). The predicted tertiary structure of TbNAB70 was generated using AlphaFold2 (66) under the AlphaFold Protein Structure Database identifier AF-Q384P8-F1.

### Conservation and synteny survey

Reciprocal BLAST searches were conducted against genes for *TbLAP1* (*Tb927.8.3060*), *TbNAB70* (*Tb927.11.7580*), *TAP110* (*Tb927.11.7590*), and *Tb927.11.6660* against a variety of publicly available Kinetoplastid, Diplonmeid, and Euglenid CDS, with a threshold E^-5^ value. Gene synteny specifically was surveyed within the genomes of Trypanosomatidae with N_50_ values of 500 kb or greater, along with *Paratrypanosoma confusum* and *Bodo saltans* (67).

## ACKNOWLEDGEMENTS

We thank Karel Harant and Pavel Talacko (Charles University, Prague) for performing LC-MS/MS analysis and Ken Stuart (Seattle Children’s Research Institute, Seattle) for the HSP70 antibody. We are grateful to Keith Gull (University of Oxford) for fruitful discussions and for the FAZ and Y1/2 antibodies. We acknowledge support from the Czech Science Foundation 22-01026S (to V.Y.) and 22-14356S (to J.L.), EU’s Operational Program “Just Transition” CZ.10.03.01/00/22_003/0000003 LERCO (to V.Y.), and Czech BioImaging grant LM2015062 (to M.T.).

## Author Contribution

Conceptualization: L.R.C., J.L.; Methodology: L.R.C., M.H., M.T., Ľ.C., M.S., I.M.D.; Validation: L.R.C., M.H.; Formal analysis: L.R.C., M.H.; Investigation: L.R.C., M.H.; Resources: J.L., V.Y.; Writing – original draft: L.R.C., M.H., J.L.; Writing – review & editing: L.R.C., M.H., V.Y., J.L.; Visualization: L.R.C., M.H.; Supervision: J.L.; Project administration: J.L.; Funding acquisition: J.L., V.Y.

