## Supplementary figures and images for "A novel nabelschnur protein regulates segregation of the kinetoplast DNA in *Trypanosoma brucei*"

### Fig. S1

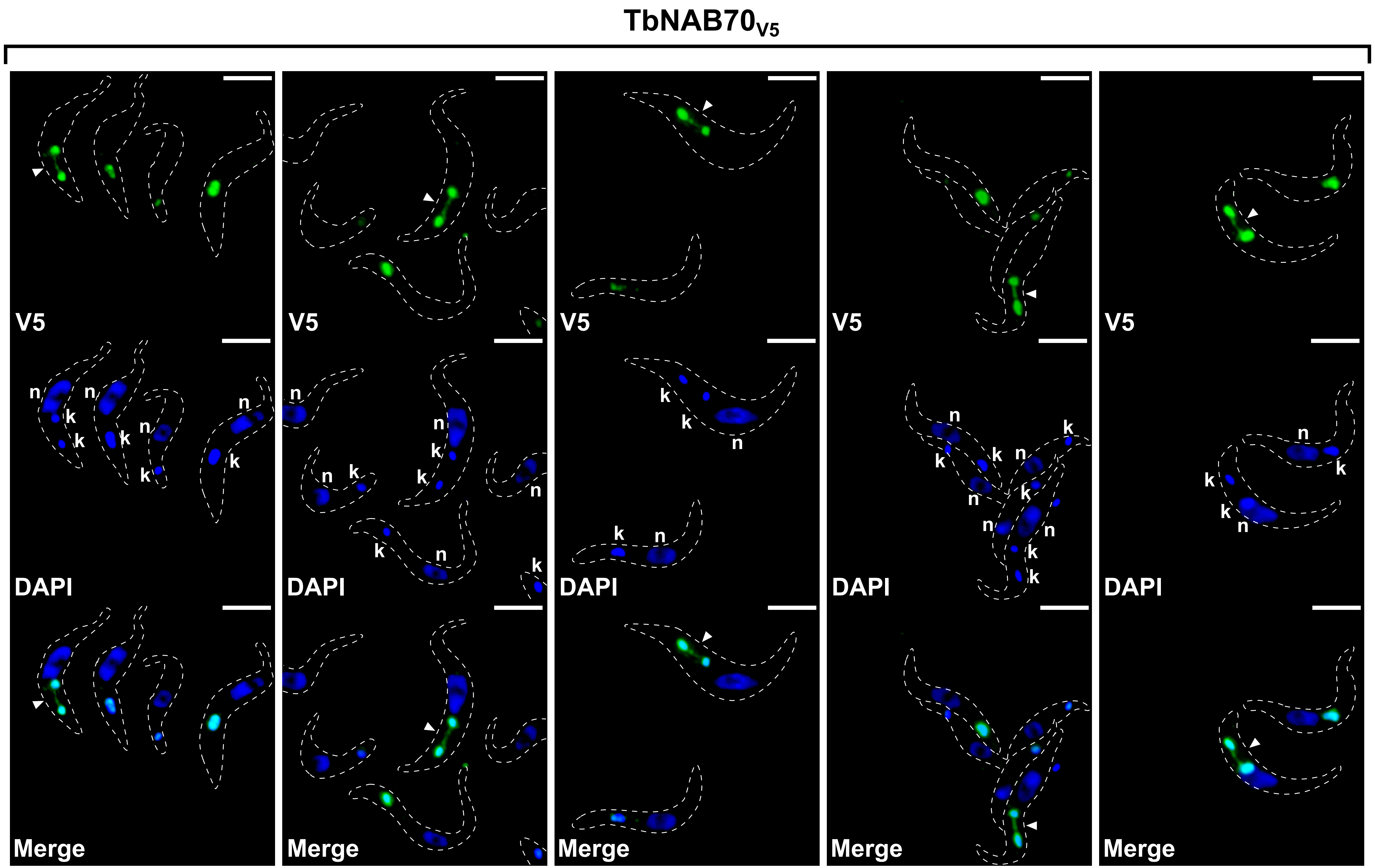
